# Oogenesis and germinal bed morphology of the brown anole (*A. sagrei*)

**DOI:** 10.1101/2025.09.11.675729

**Authors:** Bonnie K. Kircher, Antonia Weberling, Erin J. Vance, Natalia A. Shylo, Katherine Starr, Zoe B. Griffin, Hannah Wilson, Melainia McClain, Florian Hollfelder, Suzannah A. Williams, Thomas J. Sanger, Richard R. Behringer, Paul A. Trainor

## Abstract

**Background:** The brown anole is a model species of the genus *Anolis*, a squamate (encompassing lizards and snakes) group widely studied in evolutionary, behavioral, and developmental biology. Full genome annotation, the establishment of gene editing techniques, and comprehensive description of reproductive tract morphology and embryogenesis in this species, has laid the foundation for functional studies. However, analysis of brown anole oogenesis is still required and vital to optimize genome modification, mutant line establishment, and analyses of the evolution of reproductive developmental mechanisms.

**Results:** Here, we characterize ovary morphology and gametogenesis in the female brown anole, *A. sagrei* using brightfield imaging, microCT, histology staining, electron microscopy, and confocal imaging. We define 10 stages of oocyte maturation which commences inside the oogonial nest within the germinal bed and concludes with the mature follicle ready to ovulate based on follicle size, yolk-acquisition, and follicular, cellular, and basement membrane architecture.

**Conclusions:** We describe the complete oogenesis of the brown anole in 10 stages and report that oogenesis is highly conserved within iguanids, a suborder of lizards. With our staging framework, we lay the foundation for functional studies of oogenesis and optimized gene-editing.

## Introduction

Non-avian reptiles exhibit striking divergence from mammals and birds in their manner of oogenesis. In mammals and birds, oocyte development occurs during embryogenesis, pauses during meiosis I, and resumes at sexual maturity. This results in most mammals and birds containing a finite number of oocytes in their ovaries (Beaumont & Mandl, 1962; Dubois & Croisille, 1970) in contrast to non-avian reptiles (lizards, snakes, crocodilians, and turtles) (Guraya & Varma, 1976). Ovaries of lizards and snakes generate oocytes continuously from the so-called germinal bed, located at the dorsal surface of the ovary (Guraya, 2013). There, the stem cells within the oogonial nest produce primordial follicles. During this process, the oogonial stem cells transition seasonally from mitosis to meiosis, such that the stem cells in the germinal bed are mitotic while the primordial follicles transition to meiosis (Nie et al., 2023). Not all non-avian reptiles have a single germinal bed, some species can have up to two germinal beds per ovary, and each germinal bed may ovulate over 200 times per ovulatory period (Jones et al., 1982). However, despite some comprehensive histological studies of the reptilian ovary, including those in lizards, (Aldokhi et al., 2019; da Silva et al., 2018; Guraya & Varma, 1976; Klosterman, 1983; Lozano et al., 2014; Moodley & Van Wyk, 2007; Neaves, 1971; Ortiz & Morales, 1974; Uribe et al., 1995; Varma, 1970; Vieira et al., 2010) the techniques were limited to histology and ultrastructure electron microscopy across relatively few follicular stages, making comprehensive understanding of germinal bed diversity and oocyte maturation poorly understood.

In non-avian reptiles, the stages of ovarian follicle development have been described with general conservation observed between species. In short, immature follicles exit the germinal bed and are surrounded by a single layer of pre-follicular cells. As the oocyte matures, the follicle enlarges while the pre-follicular cells mature into granulosa-like cells that surround the maturing follicle (Dekel, 1987). In mammals, steroidogenic granulosa cells communicate with the oocyte on one side and with the differently steroidogenic theca cells on the other side (Nilsson & Skinner, 2001). In the non-avian reptile ovary, the granulosa-like cells are also covered by theca cells (Ramírez-Pinilla et al., 2015). As follicle development progresses, enlarged cells emerge in the granulosa cell layer including “intermediate” and “pyriform” granulosa cells. A layer of small granulosa cells lies directly over the oocyte while the enlarged intermediate and pyriform granulosa cells sit on top of the small granulosa cells, towards the outside of the follicle (Ramírez-Pinilla et al., 2015). With the emergence of these cell types, the granulosa cell layer thickens. During more mature stages of follicular development, the follicular epithelium thins, and these enlarged pyriform cells establish intracellular bridges between the follicular epithelium and the oocyte (Neaves, 1971). Although the pyriform granulosa cells are characteristic of vitellogenic follicles (Klosterman, 1983), their precise function remains unknown. At the latest, most mature stages of folliculogenesis, the epithelium becomes thinner until the mature vitellogenic, pre-ovulatory follicle is only surrounded by a single cell layer (Jones et al., 1975).

The brown anole (*A. sagrei*) has emerged as a representative species of the genus *Anolis* over the last two decades. *Anolis* comprises over 400 species that are distributed across the Caribbean, Middle and South America. Given the growing number of experimental and genomic tools, *Anolis* has emerged as a primary model for the study of evolution, ecology, behavior, and physiology (Muñoz et al., 2023). *Anolis* also serves as an important outgroup for genomic and developmental comparisons among birds, non-avian reptiles, and mammals (Gredler et al., 2014; Griffing et al., 2022; Malkmus et al., 2021; Marchini et al., 2025). Numerous tools have laid the basis for functional studies in for the brown anole, most notably a high-quality annotated genome (Geneva et al., 2022), the establishment of CRISPR/Cas9 gene edited knock outs (Rasys et al., 2019) and the generation of immortalized fibroblast lines (Samudra et al., 2024). However, CRISPR-Cas9 genome editing requires injection of pre-ovulatory follicles and currently has a success-rate of only 10-15% (Rasys et al., 2019). A better understanding of reproductive and developmental progression is needed to increase the efficiency of these techniques and for further functional studies.

The morphology of the reproductive tract and embryogenesis have been described for the brown anole (Kircher et al., 2023; Sanger et al., 2008; Weberling et al., 2025). Brown anole females have a duplex reproductive tract with uteri that coalesce at the cloaca (Kircher et al., 2023). Ovulation occurs every 5-10 days alternating between the right and the left ovary into the glandular uterus (Crews, 1977, 1980) resulting in about one egg being laid per week during the breeding season from May-September.

Oogenesis has previously been studied in two iguanid genera. In *Anolis pulchellus*, 4 stages of follicular maturation have been described (Ortiz & Morales, 1974) that loosely correlate with the 9 more detailed stages defined for four *Tropidurus* species (da Silva et al., 2018). However, a detailed description of ovary and follicle development in the genus *Anolis* does not exist. To enable higher success rates for gene editing and inform functional studies, we need to better understand ovarian follicular morphology, development, and maturation.

Here, we provide a detailed description of *Anolis sagrei* oogenesis and ovarian morphology using brightfield imaging, computed tomography (CT)-scans, scanning transmission electron microscopy (STEM), and histology staining as well as confocal imaging. Guided by the description of follicle development in 9 stages in *Tropidurus* (da Silva et al., 2018), we define 10 consecutive stages of oogenesis in the brown anole, including one novel stage preceding the stages described in *Tropidurus*. We analyze follicle development through assessment of overall follicle size, the width and architecture of the follicular epithelium, and the basement membrane composition. Taken together, our study lays the foundation for functional studies of non-avian reptile oogenesis and optimized CRISPR/Cas9 genome editing.

## Results

### Brightfield and volumetric analysis of ovarian follicle growth

The ovary of the brown anole is composed of follicles that vary significantly in size and color (Figure 1A-C). The smallest follicles are colorless, then they become white and eventually beige due to yolk acquisition while increasing in size. These immature follicles are found in varying numbers in every ovary. In addition to these follicles, each ovary contains a maximum of one mature, yellow yolk-filled follicle, which is much larger than the beige follicles. It exhibits a bright yellow color and is enveloped by a vascularized, thin tissue (Figure 1B/D, asterisks). The germinal vesicle can be observed in every follicle (Figure 1A, higher magnification) but becomes more apparent upon the initiation of vitellogenesis (Figure 1C asterisk). In mature follicles, the germinal vesicle is found in the center of a white plate, the germinal disk (Figure 1C, dashed lines, Figure 1C/D arrows). To quantify the dramatic size increase between immature, maturing, and the mature follicle and to understand whether these characteristic size differences were also present *in situ*, we assessed the volume of ovarian follicles following microCT imaging of whole females (Figure 1E). The ovaries are located dorsally in the coelomic cavity of the lizard, just anterior to the cloaca (Figure 1E). We digitally segmented the follicles on both the left and right ovaries (Figure 1, Figure 2), where we observed the increase in size between the immature (yellow/cyan/green highlight) and the mature follicles (pink highlight) (Figure 1F-G). Plotting the follicle volume, we hypothesized that ovarian follicle size increases exponentially (Figure 1H). Following quantitative analysis of left and right ovaries of three females, we confirmed an exponential increase in follicle volume occurred during oogenesis (Figure 1I, Table 2). This trend also holds true when analyzing follicle volume across all three biological replicates at once (Figure 1I; p= 6.656e-10, R^2^= 0.6483, F(1,36)= 69.2). We did not observe significant differences between left and right ovaries and therefore the data were pooled for further analysis (individual statistics in Table 1). Taken together, our volumetric analysis of different-aged follicles in anole ovaries revealed an exponential volume increase during maturation.

**Figure 1:**
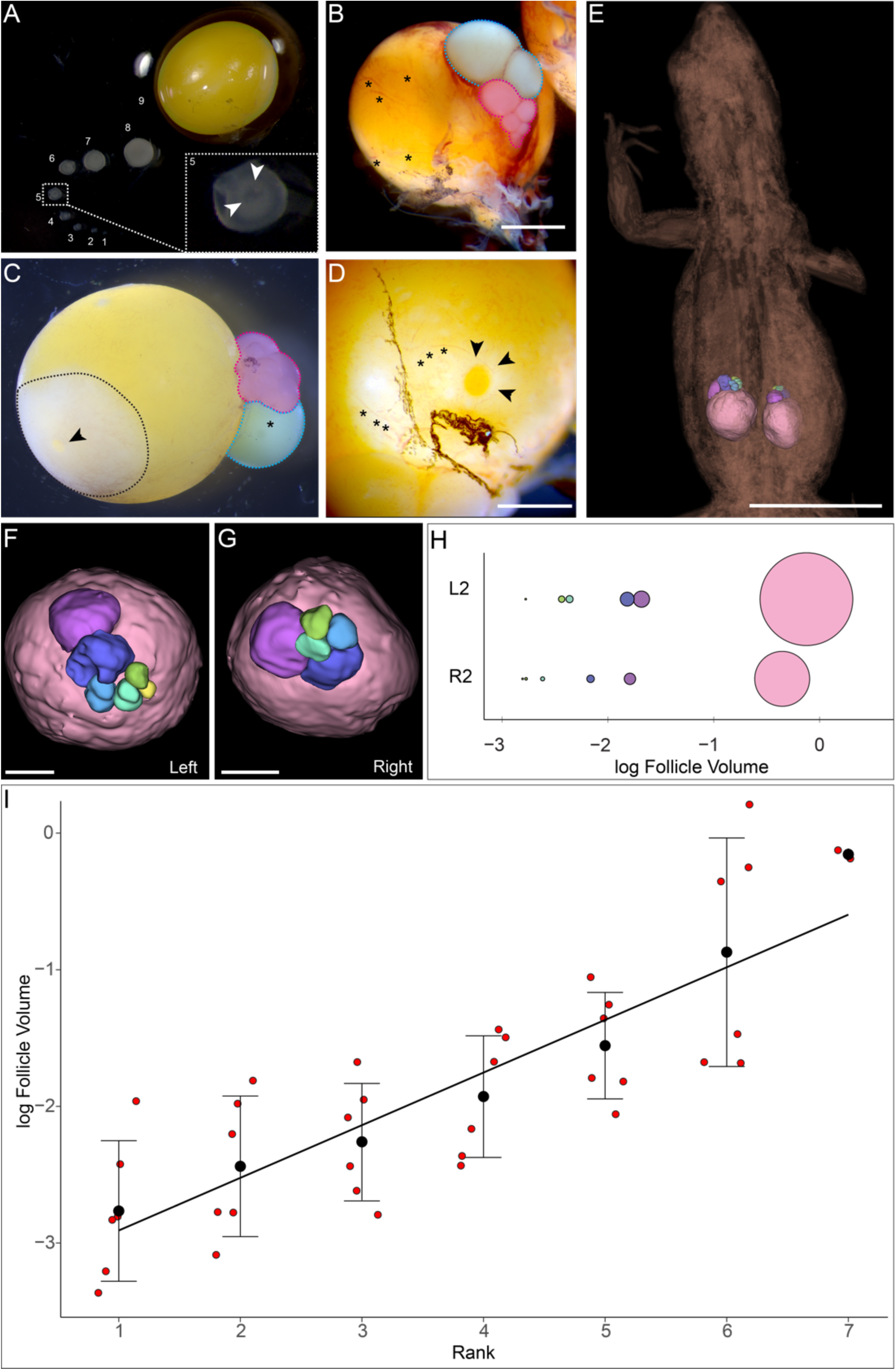
Volumetric analysis of the brown anole ovary A-D. Brightfield images of anole ovaries. (A) 9 follicles dissected from a single ovary with (1) as smallest and (9) as largest. Higher magnification of follicle (5), white arrows indicate the germinal vesicle. Ovary morphology (B/C) early follicles (magenta highlight), initiation of yolk acquisition (cyan highlight), (B) asterisks blood vessels, scale bar = 2mm. (C) asterisk in cyan highlighted follicle germinal vesicle. Dashed outline: germinal disk, arrow germinal vesicle. (D) higher magnification of the germinal disk. Asterisks: blood vessels. Arrows highlight the germinal vesicle, scale bar = 1mm. **E-G.** CT scans of ovaries from fixed anole females. Body visible in brown. Individual follicles segmented and color-coded. (E) position of ovaries within the abdomen, scale bar = 10mm. (F) higher magnification of the left ovary, scale bar = 1mm. (G) higher magnification of the right ovary, scale bar = 1mm. **H.** volumetric analysis of follicle size in left and right ovaries. Logarithmic scale. **I.** Quantitative analysis of follicle volumes of six ovaries. Scatter plot mean±StDev. Rank is assigned based on follicle size so that the smaller follicles have a lower rank.

**Figure 2:**
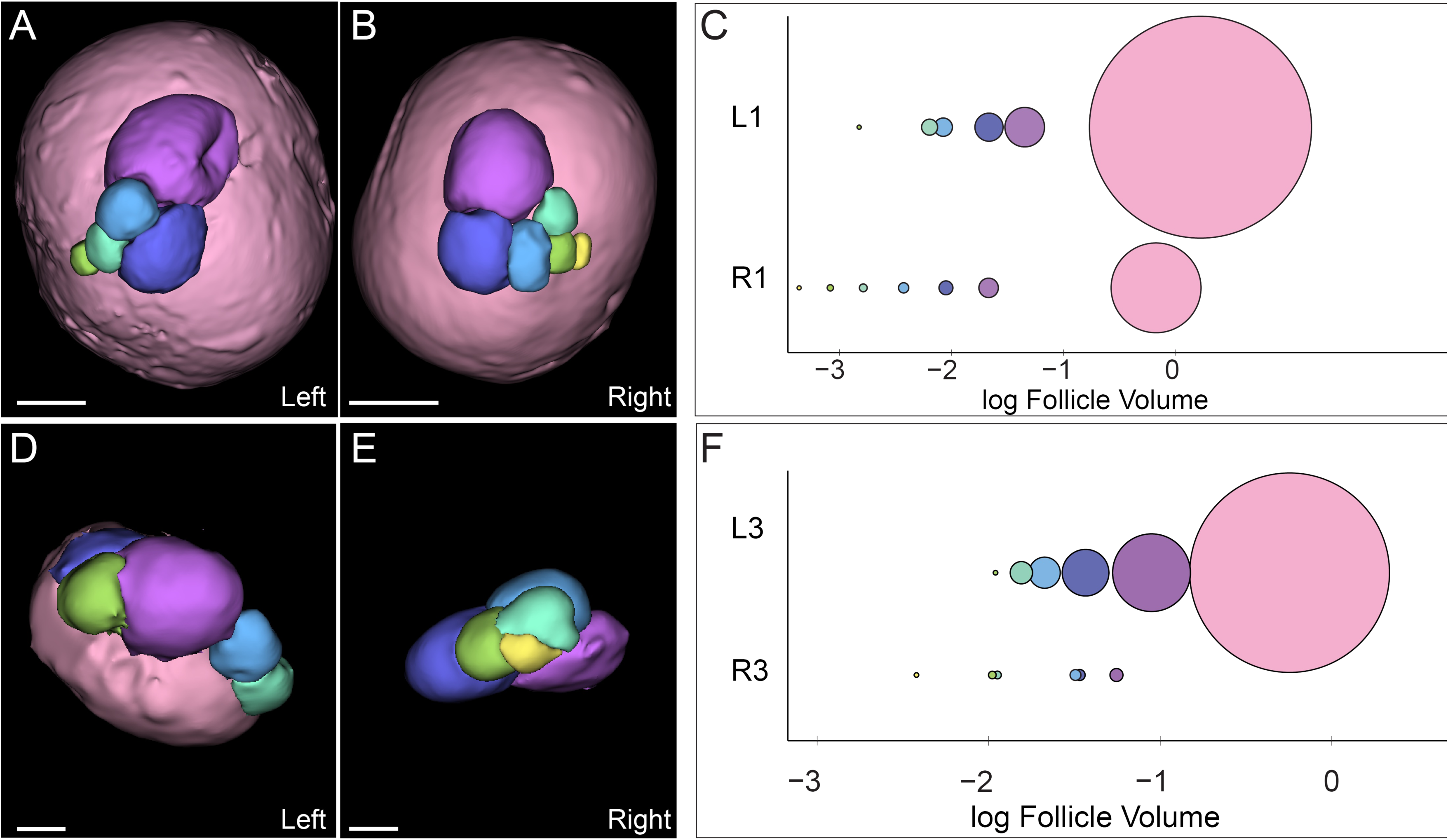
CT scans of brown anole ovaries A/B. CT scans of ovaries from fixated anole females. Individual follicles segmented and color-coded. Replicate 2 (Anole 1) with higher magnification of the left ovary (A) and higher magnification of the right ovary (B), scale bar = 1mm. Replicate 3 (Anole 3) with higher magnification of the left ovary (D) and higher magnification of the right ovary (E), scale bar = 1mm. **C,F.** Volumetric analysis of follicle size in left and right ovaries.

**Table 1.**
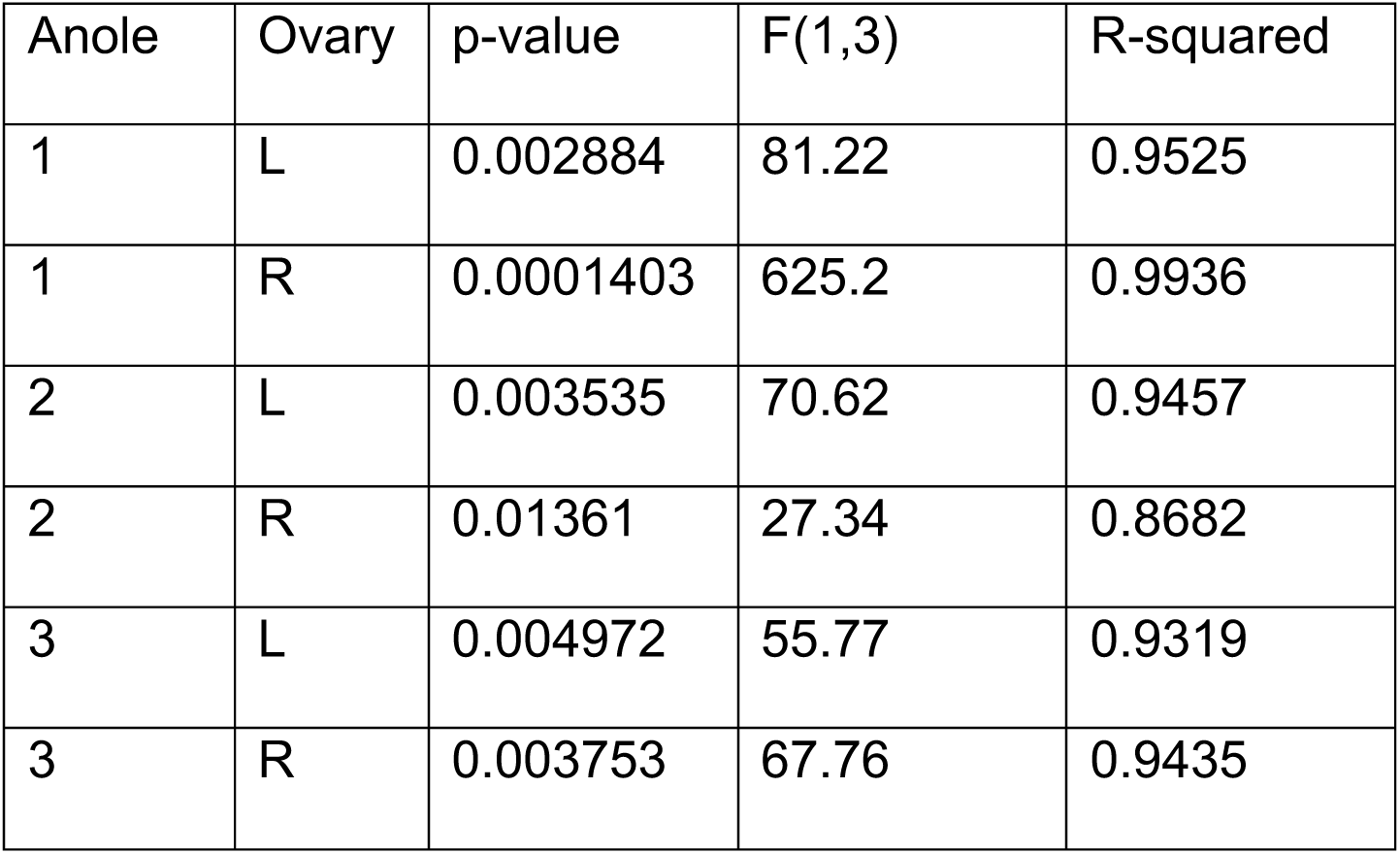
Regression statistics for follicular volume across six individual ovaries analyzed.

**Table 2.**
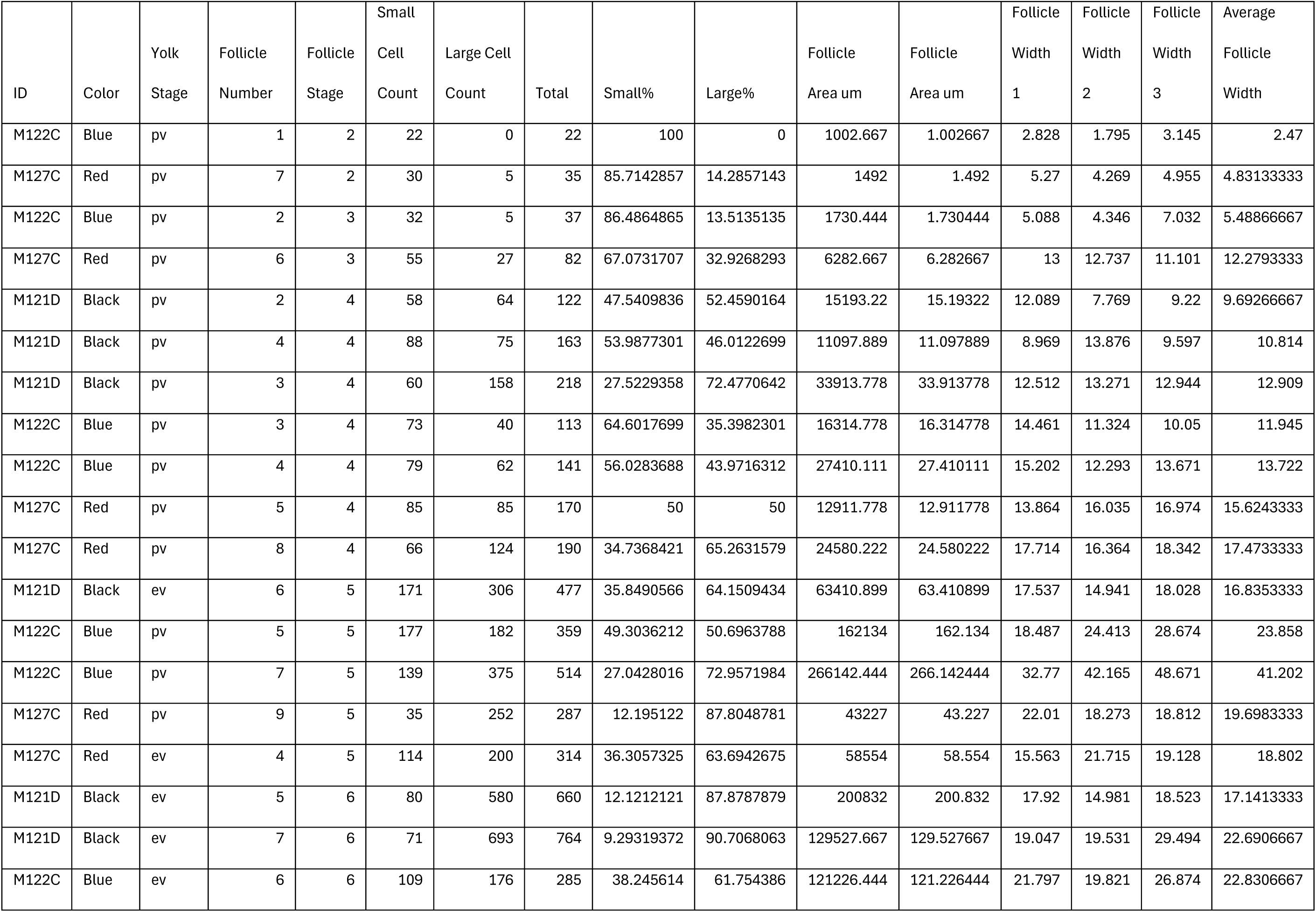
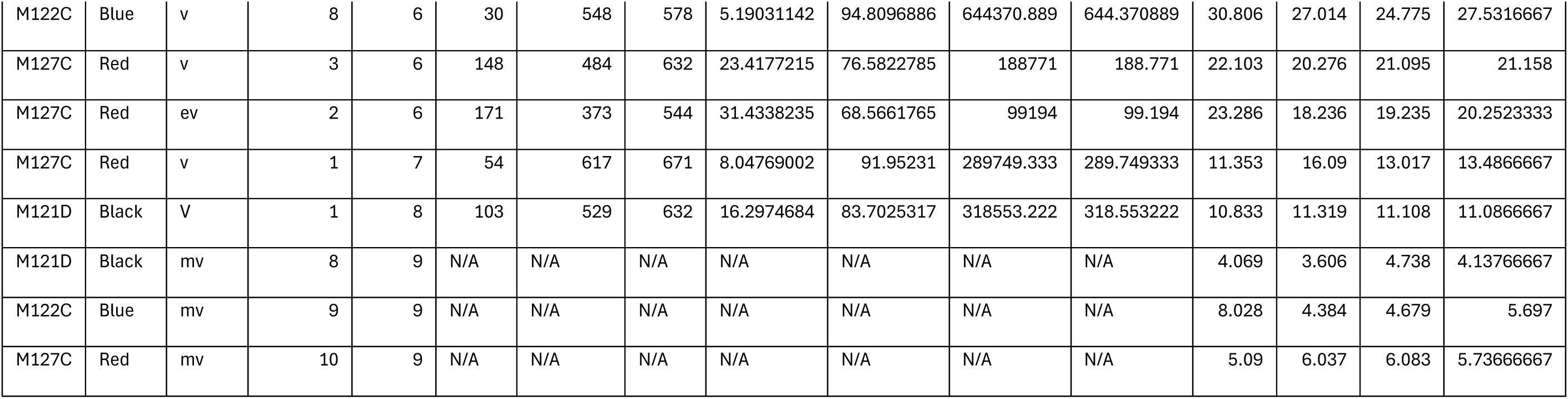

### Follicle maturation is accompanied by changes in yolk density patterns

Following our brightfield microscopy assessment that suggested progressive yolk acquisition upon follicle maturation, we collected cross-sections of whole ovaries and individual germinal vesicles and analyzed them using hematoxylin and eosin (H&E) staining. H&E staining of whole ovary sections revealed distinct density patterns within the yolk of different sized follicles (Figure 3A-E, Figure 4). Mature vitellogenic follicles are composed of homogenous, dense yolk that has a grainy appearance (Figure 3A, arrows, 3B/B’) surrounded by a thin cell monolayer (Figure 3B arrows). Maturing vitellogenic follicles exhibit three distinct layers of yolk (Figure 3C/C’/C’’). The outer-most layer 1 is dense and rich in lipid droplets (Figure 3C’ arrows). These characteristics are in stark contrast to layer 2, which appears spongy with a low density, and fills most of the follicle volume (Figure 3C/C’/C’’ 2). In the middle of the follicle, a small third layer can be found, which again exhibits higher density, but no large lipid droplets as observed in layer 1 (Figure 3C/C’’ 3). Early vitellogenic follicles are composed of 2 different layers (Figure 3D/D’/D’’). The outer layer, which constitutes most of the follicle, is dense and homogenous (Figure 3D/D’ 1). The inner layer appears loose and spongy, like layer 2 in the maturing vitellogenic follicle (Figure 3D/D’’). In pre-vitellogenic follicles (Figure 3E), the yolk/cytoplasm has a homogenous structure and only exhibits small foci of high density surrounded by lower density (Figure 3E’ asterisks). Interestingly, we observed a set of structures inside the germinal vesicle that may be condensed DNA (Figure 3E’’ arrows). To understand these qualitative observations in more detail, we measured yolk density by quantifying the changes in gray-value across the histological sections of the four stages of yolk development using plot profiles (Figure 3F, Figure 4). Gray values are higher and more consistent in pre-vitellogenic follicles correlating with the homogeneous structure of the yolk/cytoplasm (Figure 3F). In early and vitellogenic follicles, the gray values are higher in the middle of the follicle compared to the outer edges of the follicle, correlating with the layered morphology of the yolk developing at these stages (Figure 3F). In mature vitellogenic follicles, gray values are variable across the plot profile, reflecting the homogeneous lipid droplet architecture in the cytoplasm of these follicles.

**Figure 3:**
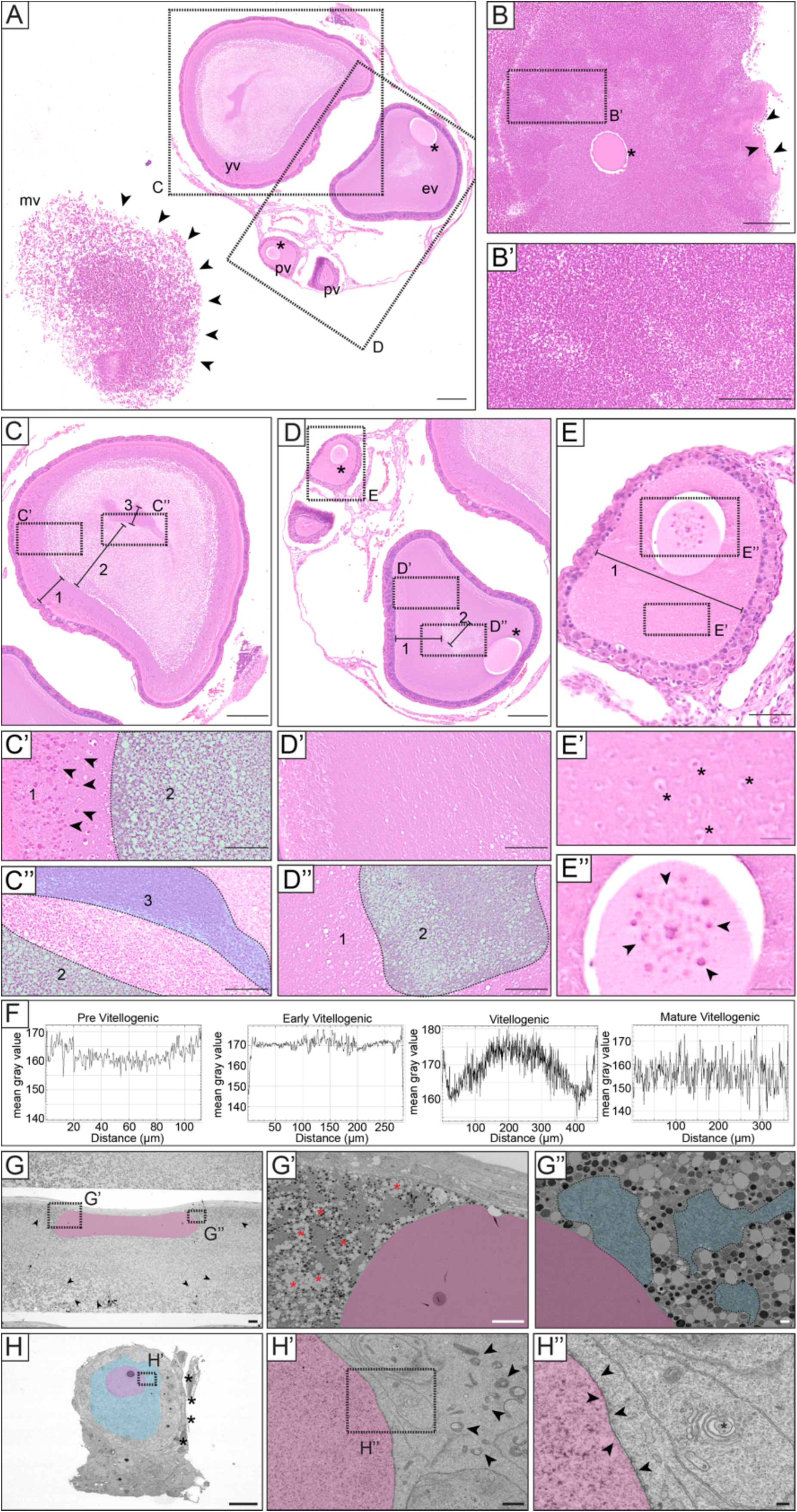
Analysis of yolk composition during vitellogenesis. **A.** H&E staining of ovary cross section. The ovary contains a vitellogenic follicle (mv). A maturing vitellogenic follicle (yv), an early vitellogenic follicle (ev) and several previtellogenic follicles (pv). Arrows indicate the mature vitellogenic follicle that broke apart during processing, box C indicates the higher magnification of (C), box D indicates higher magnification of (D), scale bar = 200um. **B.** H&E staining of mature vitellogenic follicle. Asterisk indicates germinal vesicle arrows indicate cell layer surrounding follicle. Box B’ outlines higher magnification of (B’), scale bar 200um. **B’.** higher magnification of yolk in mature vitellogenic follicle. Yolk dense and homogenous, scale bar = 100um. **C.** H&E staining of maturing vitellogenic follicle. Yolk exhibits 3 (1-3) distinct layers of different density and composition, scale bar = 200um **C’.** higher magnification oflayer 1&2. Layer 1 exhibits large darker droplets (arrows). Layer 2 (green highlight) is less dense, scale bar = 50um. **C’’.** Higher magnification of layers 2 (green) and 3 (cyan), which is denser than 2, scale bar = 50um. **D.** Early vitellogenic follicle. Yolk is composed of 2 layers (1-2) boxes D’/D’’ indicate higher magnifications. Asterisks indicate germinal vesicles. Box E indicates higher magnification of (E), scale bar = 200um. **D’.** homogenous yolk composition in layer 1, scale bar = 50um. **D’’.** Yolk composition of layer 1 versus layer 2 (green highlight), scale bar = 50um. **E.** Pre-vitellogenic follicle. Yolk is homogenous (1). Boxes indicate higher magnifications, scale bar 50um. **E’.** Yolk composition of pre-vitellogenic follicle. Asterisks indicate small foci of less density, scale bar = 10um. **E’’.** Germinal vesicle exhibits dense foci (arrows) that could be DNA, scale bar 20um. **F.** Plot profiles of yolk composition of pre-vitellogenic, early vitellogenic, vitellogenic and mature vitellogenic follicles. **G.** STEM of mature vitellogenic follicle. Germinal vesicle highlighted in magenta. Yolk is filled with droplets (arrows). Boxes indicate higher magnifications (G’/G’’). scale bar = 20um. **G’.** The yolk is filled with lipid droplets (asterisks), scale bar = 10um. **G’’.** High concentrations of mitochondria are observed (blue highlight), scale bar = 1um. **H.** STEM of previtellogenic follicle. Germinal vesicle highlighted in magenta. Cytoplasm/yolk (blue highlight) lacks lipid droplets and the punctae-like structure found in (G), box indicates higher magnification (H’). asterisks indicate germinal bed, scale bar = 20um. **H’.** Cytoplasm/yolk exhibits low density of mitochondria and no lipid droplets. scale bar = 1um. **H’’.** Double membrane (arrows) surrounding the germinal vesicle not in direct proximity of lipid droplets or mitochondria, asterisk indicates golgi apparatus, scale bar = 200nm.

**Figure 4:**
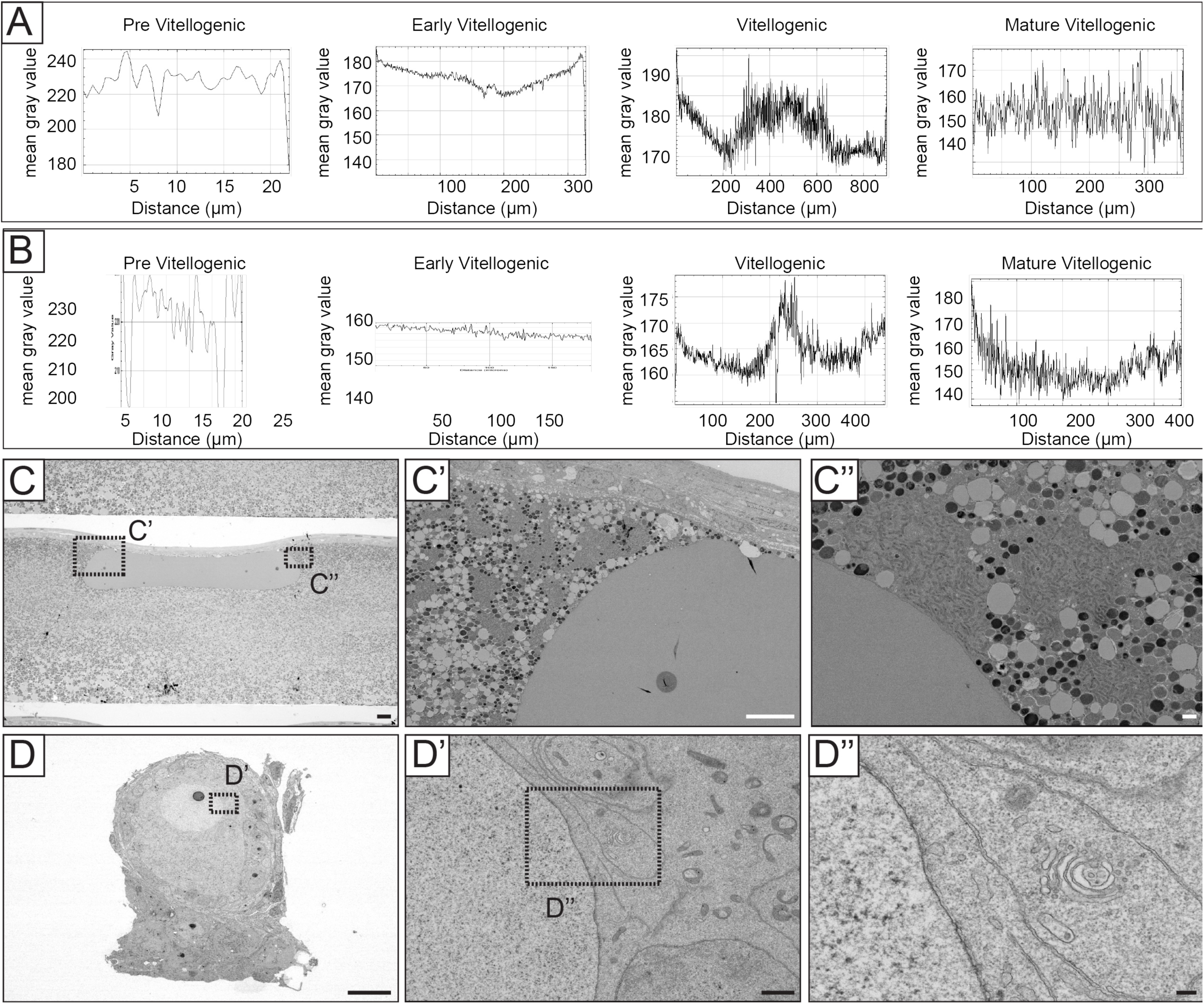
Analysis of yolk composition during follicle maturation A.-B. Plot profiles of mean gray value of yolk composition during follicle development. Biological replicates of Figure 2F. **C.-D.** STEM of previtellogenic and mature vitellogenic follicle. Images from F2G/H without annotations and highlights. Scale bar for C = 20um, scale bar for D = 200nm.

To gain higher resolution insight into the substructure of the yolk in vitellogenic versus previtellogenic follicles, we performed Transmission Electron Microscopy (TEM) (Figure 3G-H, Figure 4). The yolk of vitellogenic follicles is rich in lipid droplets (Figure 3G arrows, 3G’ asterisks) and exhibits high numbers of tightly packed mitochondria (blue highlight) in direct proximity to the germinal vesicle (magenta) (Figure 3G’’). In contrast, pre-vitellogenic follicles do not contain lipid droplets in their cytoplasm (Figure 3H, blue highlight). Mitochondria can be observed but at much lower density than in vitellogenic follicles (Figure 3H’, arrows) and are not found in direct proximity to the germinal vesicle (Figure 3H’’). Instead, elongated ER and Golgi apparatus (asterisk) were observed surrounding the germinal vesicle (Figure 3H’’). Interestingly, we identified the germinal bed in direct proximity to the pre-vitellogenic follicle (Figure 3H asterisks).

### The different stages of oogenesis can be distinguished through the changes in the follicular epithelium

Following our observation of the germinal bed (Figure 3H), we examined the progression of ovarian follicle development and maturation from oogonia to a mature vitellogenic follicle. Therefore, we carried out H&E staining of cross-sections of ovaries (Figure 5A-C, Figure 6). In the germinal bed, we observed oogonia and primordial follicles (Figure 5A). Upon initiation of follicle development, we observed distinct differences in the follicular epithelium. Across all follicle stages, epithelium thickness ranged from a single cell layer in the earliest stages (Figure 5A blue highlight) to a thick, multilayered envelope of different cell types in the later stage follicles (Figure 5B). The mature vitellogenic follicle exhibits again only a single cell layer in the mature vitellogenic follicle (Figure 5C). We then analyzed follicle morphology and composition of the follicular epithelium (Figure 5D, 6D). In the germinal bed, two distinct populations of germ cells can be observed. Numerous oogonia cluster into the oogonial nest (Figure 5D1, blue highlight). The individual oogonia are characterized by small, dense nuclei and cytoplasm devoid of eosin staining. The nuclei are not round but rather exhibit signs of mitosis, such as aligned or condensed chromosomes (Figure 6D1i). We term this **stage Ia**. The second population we could define in the germinal bed are primordial follicles (Figure 5D1, yellow highlight 6D1ii). Here, the nuclei are no longer condensed but round and of lighter staining. The diameter of these cells has enlarged by a factor of two-three compared to the oogonia, and the cytoplasm is positive for eosin. These primordial follicles are not as tightly packed as the cells in the oogonial nest. Instead, they appear interspersed with epithelial cells. We termed this **stage Ib**.

**Figure 5:**
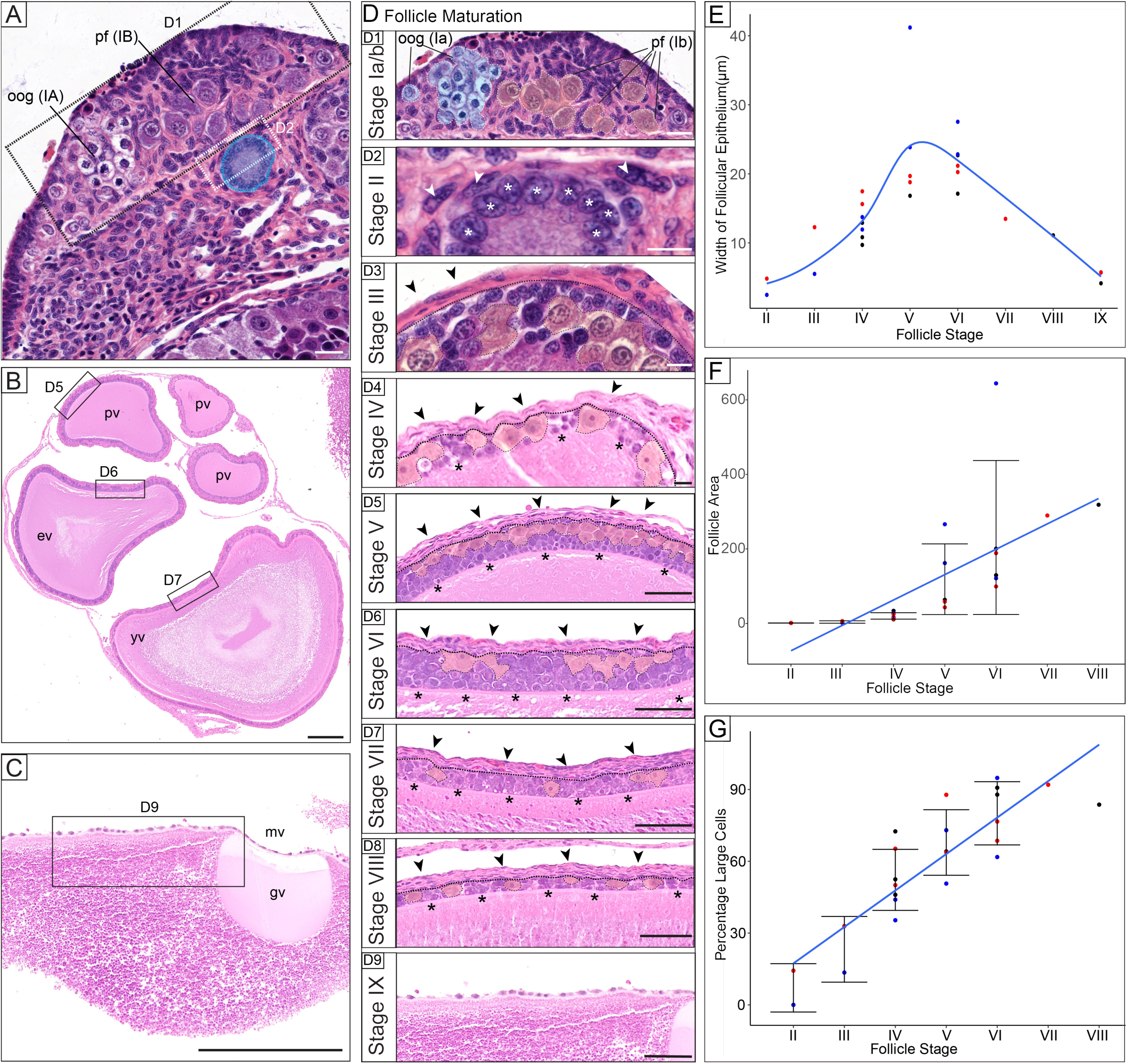
Histological analysis of germ cell maturation. **A.** H&E staining of ovarian cross section containing the germinal bed populated with oogonia (oog) and primordial follicles (pf), and a stage II follicle (blue highlight). Rectangles indicate higher magnification of Figure D1/D2. scale bar = 20um. **B.** H&E staining ovarian cross section of previtellogenic to vitellogenic follicles. pv = previtellogenic, ev = early vitellogenic, yv = yolk vitellogenic. Rectangles indicate higher magnification of Figure D5-7. scale bar = 200um. **C.** H&E staining cross section mature vitellogenic follicle (mv). Germinal vesicle (gv) visible. Square indicated higher magnification of Figure D9. scale bar = 200um. **D.** H&E staining of the 9 stages of follicle maturation (I-IX). (D1) oogonia (blue highlight, oog), primordial follicles (yellow highlight, pf), scale bar = 20um, (D2-9) granulosa cells (asterisks) and theca cells (arrows) are highlighted, D2-4 scale bar = 10um, D5-9 scale bar = 50um. (D4-8) dashed line between theca and granulosa cell layer, pyriform granulosa cells highlighted in yellow. all scale bars = 20um. **E.** Quantitative analysis of the thickness of the follicular epithelium (granulosa & theca cells combined) from stage I-IX. **F.** Quantitative analysis of follicle size from stage Ia-stage IX. **G.** Quantitative analysis of the granulosa cell layer. The percentage of large (pyriform) granulosa cells plotted over stages I-VIII. Ovaries M121D (black), M127C (red), M122C (blue) were analyzed. Trendline is a linear model in R. Adjusted Statistical analysis: R-squared: 0.7605, F-statistic: 74.01 on 1 and 22 DF, p-value: 1.73e-08.

**Figure 6:**
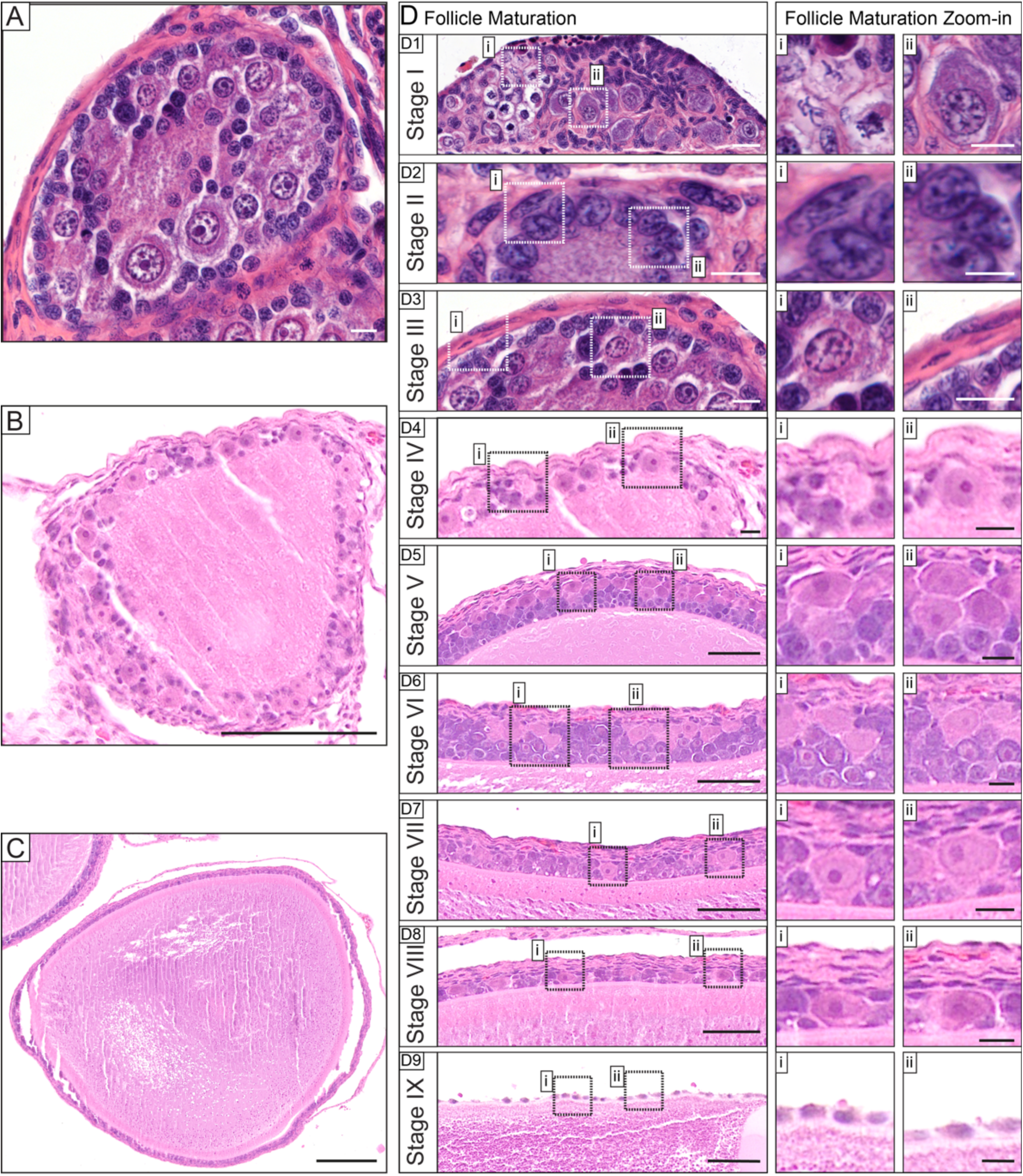
Histology of the consecutive stages of oocyte maturation. **A.** H&E staining of stage III follicle, higher magnification shown in Figure 3D3. Scale bar = 10um. **B.** H&E staining of stage IV follicle, higher magnification shown in Figure 3D4. Scale bar = 100um. **C.** H&E staining of stage VIII follicle, higher magnification shown in Figure 3D8. Scale bar = 200um. **D.** H&E staining of the stages of anole follicle development, non-annotated copy of Figure 3D. higher magnifications illustrate different stage characteristics. D2-4 scale bar = 10um, D5-9 scale bar = 50um. D1-2, i-ii scale bar = 5um, D3-9, i-ii scale bar = 10um.

At the next stage (**stage II**), the follicle size increases further and is enveloped by a tightly packed layer of granulosa cells surrounding the oocyte (Figure 52). The granulosa cells exhibit characteristic rounded nuclei (asterisks). We also observed the first theca cell layers surrounding the granulosa cells defined by their elongated nuclear shapes (arrows).

At **stage III** (Figure 53), the follicle has a diameter of 4±2.3 mm (Supplementary table 1). The first pyriform granulosa cells (yellow highlight) appear in a multilayered granulosa layer that does not exhibit clear organization of the different cell types. The theca cells (arrows) have formed a double to triple-layer on top of the granulosa cells.

At **stage IV**, as the follicle continues to enlarge (Figure 54), the number of pyriform granulosa cells increases while the intermediate (yellow highlight) and small granulosa cells are localized closer to the follicle (asterisks). The theca cells continue to be arranged in a minimum of three layers (Figure 54, arrows).

At **stage V**, the granulosa cell layer has matured into multiple layers of small and intermediate granulosa cells close to the follicle (Figure 55, asterisks), while the pyriform granulosa cells (yellow highlight) form a mono-to double layer on top of the small and intermediate cells (asterisks). The theca cells have increased to 6-7 layers (arrows).

At **stage VI**, the follicle has initiated vitellogenesis (Figure 5B ev), and the pyriform granulosa cells no longer form a continuous layer (Figure 56, yellow highlight). The remaining granulosa cell layer is composed almost exclusively of intermediate cells (asterisks), with very few remaining small granulosa cells.

At **stage VII**, the granulosa cell layer has significantly thinned, decreasing from ∼5 to 2-3 cell layers (Figure 57, 6D7). Pyriform granulosa cells contact the oocyte (yellow highlight). As no small, and only a few intermediate granulosa cells can be observed, most of the granulosa layer is populated by large granulosa cells. The theca cell layer remains unchanged at this stage.

At **stage VIII**, the granulosa cell layer has thinned to 1-2 cell layers (Figure 58). All pyriform granulosa cells (yellow highlight) are now in direct contact with the oocyte and have adopted a squamous shape. The theca cell layer remains unchanged which results in about 50% of the follicle lining being composed of theca cells.

The mature follicle at **stage IX** has lost the granulosa cell layer and is covered by a thin thecal cell layer that surrounds the entire follicle (Figure 59, 6D9).

We quantified changes in the follicular epithelium width across the different follicle stages described above. Upon the initiation of vitellogenesis, the width of the epithelium increases, but then, following an abrupt thinning of the follicular epithelium at stage VII, it returns to a single cell layer in mature, pre-ovulatory follicles (Figure 5E). We then assessed follicle area across the different stages noting an increase as follicles mature (Figure 5F). Finally, we measured changes in the cell composition of the granulosa layer by counting small and large (including both intermediate and pyriform) granulosa cells. We observed the population of large granulosa cells increases over time while the small granulosa cells gradually disappear (Figure 5G).

### The subcellular structure of stage I-II and stage VIII follicles

Next, we focused on the subcellular structures at the initiation of stage II, and during later, stage VIII, follicle development using TEM (Figure 7/8). We therefore dissected individual follicles and studied the smallest follicle (Figure 7A/8A). Following our stage descriptions (Figure 5D), we defined the follicle as stage II, as it exhibits a single layer of granulosa cells, which have oval nuclei (blue highlight), and a thin layer of theca cells which present with highly elongated nuclei (green highlight) (Figure 7A’/8A’). On the right-hand side of the follicle, we observed a group of cells with enlarged, more rounded nuclei and prominent nucleoli reminiscent of the structure of the germinal vesicle inside the follicle (Figure 7A’’, 8A’’). We hypothesize that these are the oogonia based on their size (Grier et al., 2016) and our comparison to the oogonial nest described above (Figure 5D1). Beneath the follicle, we observed a group of 4 cells exhibiting markedly enlarged nuclei and cytoplasm (Figure 7A’’’/8’’’). We hypothesize that these cells are primordial follicles as they do not have organized granulosa cells around them but are interspersed with cells that may mature into the granulosa layer.

**Figure 7:**
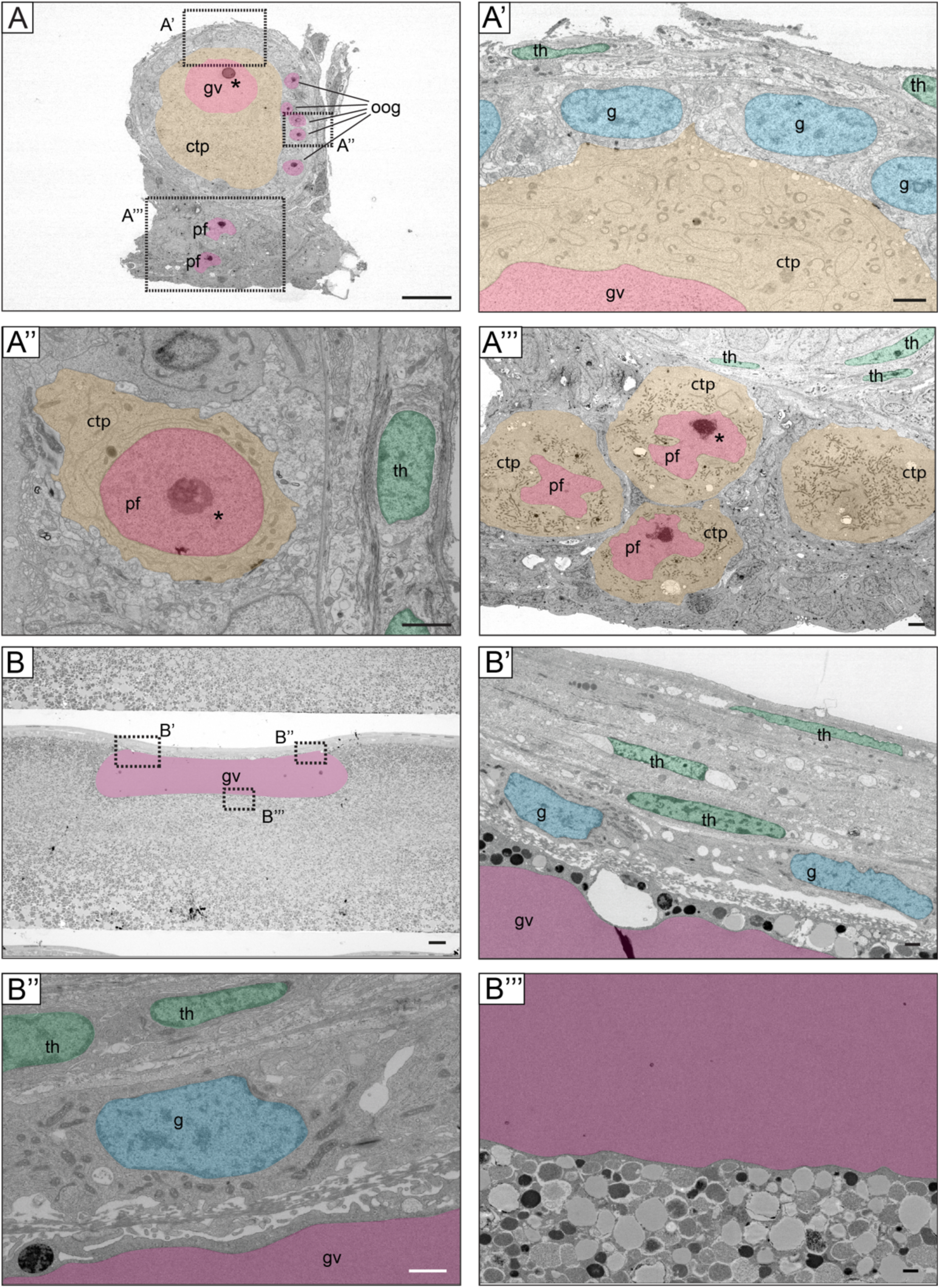
Subcellular architecture of the germinal bed A/B. STEM of germinal bed and stage II follicle (A and stage VIII follicle (B). squares indicate regions for higher magnifications. Color code: germinal vesicle/germ cell nuclei (magenta highlight), cytoplasm of primordial/previtellogenic follicles (yellow highlight) granulosa cell nuclei (blue highlight), theca cell nuclei (green highlight), asterisks indicate nucleoli. ctp = cytoplasm, g = granulosa cell, gv = germinal vesicle, oog = oogonia, pf = primordial follicle, th = theca cell. Scale bars: A/B = 20um. A’-A’’’ = 2um, B’-B’’’ = 1um

**Figure 8:**
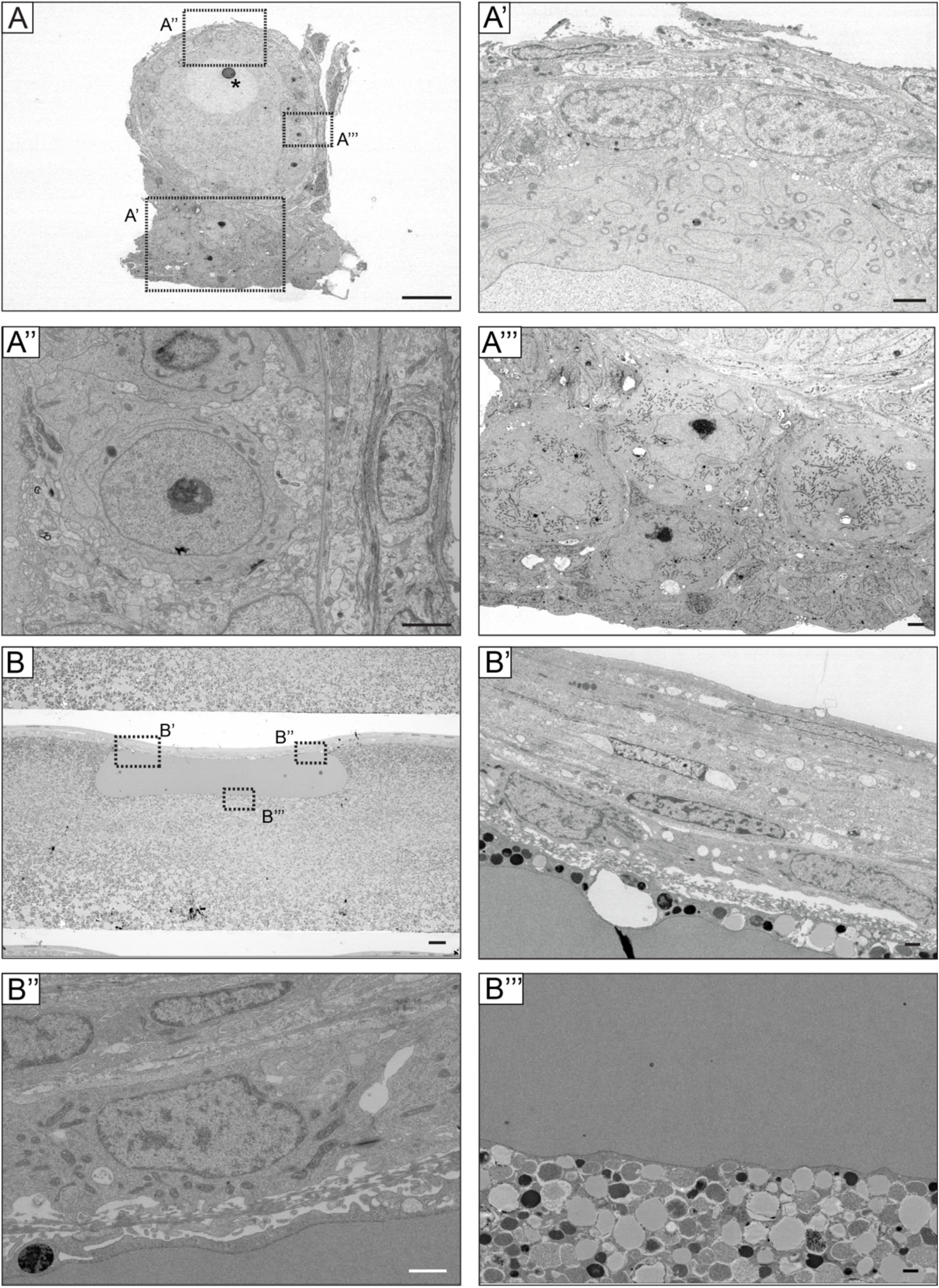
Subcellular architecture of the germinal bed II A/B. STEM of germinal bed and stage II follicle (A and stage VIII follicle (B). non-annotated images shown in Figure 4. squares indicate regions for higher magnifications. Scale bars: A/B = 20um. A’-A’’’ = 2um, B’-B’’ = 1um

We then focused on a stage VIII follicle (Figure 7B/8B) and observed a monolayer of granulosa cells overlaying the follicle (Figure 7B’/’’, 8B’/’’). The nuclei of these granulosa cells adopted a squamous shape (blue) confirming our observations made with H&E staining (Figure 5). The theca cell nuclei are highly squamous (green highlight), and the theca cell layer is about two-thirds of the thickness of the follicular epithelium. Finally, we compared the subcellular architecture of the cytoplasm/yolk of stage II/stage VIII follicles, and found that at stage II, the cytoplasm is populated by organelles such as endoplasmic reticulum and mitochondria, while the yolk of the stage VIII follicle is filled with lipid droplets (Figure 7A/8A, 7B’’’/8B’’’). Our results detailed and confirmed characteristic changes in the follicular epithelium upon follicle maturation.

### Basement membrane architecture during follicle development

To investigate changes in basement membrane composition and thickness during follicle maturation, we performed trichrome staining on cross sections of ovaries (Figure 9A-E). The germinal bed was separated from maturing follicles by a thick layer of basement membrane (asterisks) (Figure 9A/A’ asterisks). Upon initiation of follicle development, the follicles appear enveloped in a thick layer of basement membrane (Figure 9A’ arrows). In stage III follicles (Figure 9A’’), we observed a very thin basement membrane layer around the oocyte and a thick layer of basement membrane in the theca cell layer.

**Figure 9:**
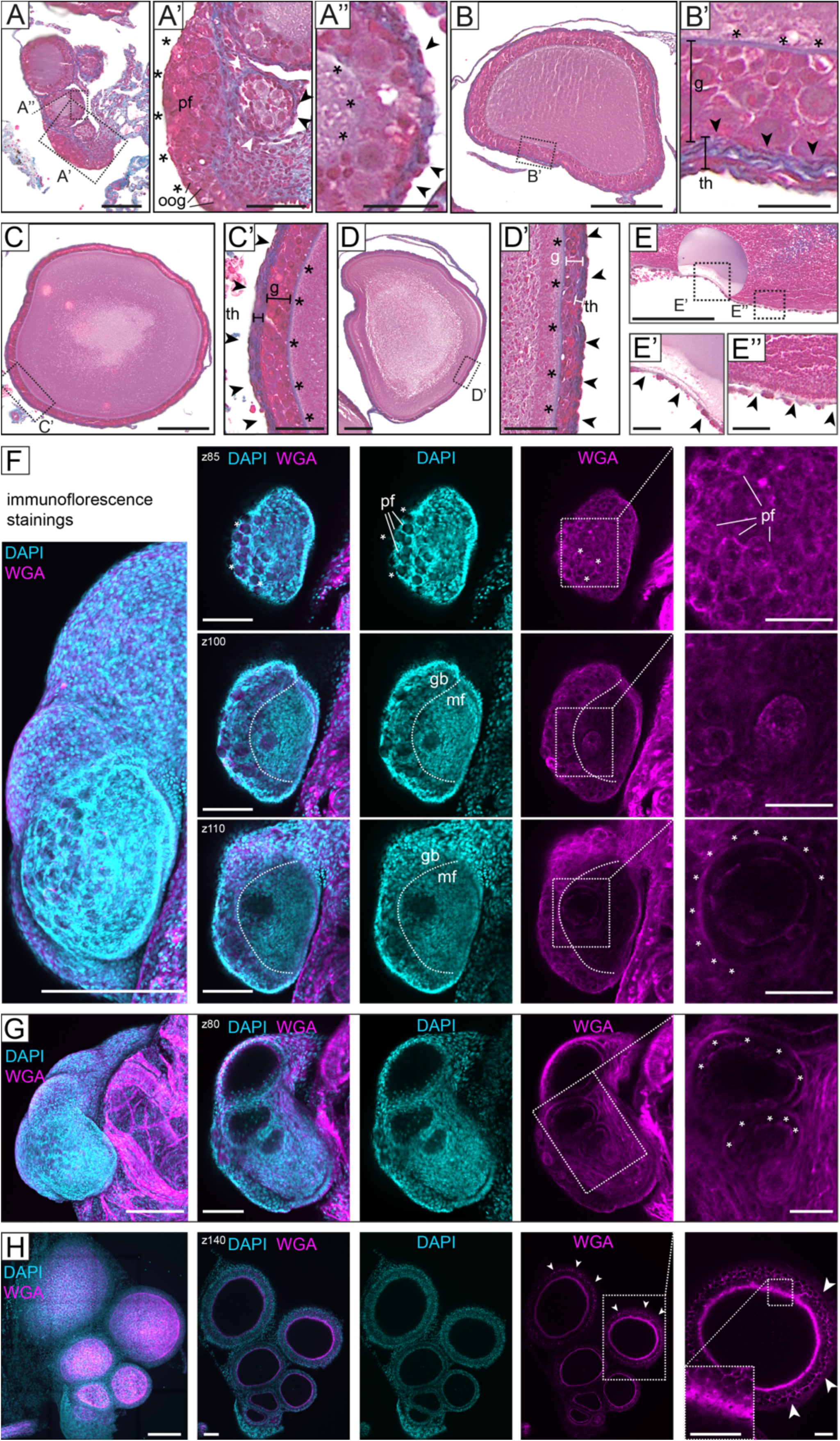
Basement membrane deposition during follicle maturation A-E. Trichrome staining of stage I-IX follicles on cross sections. Square indicates higher magnifications. (A) stage I-II, scale bar = 100um (A’) higher magnification of oogonia (oog, asterisks) and primordial follicles (pf, arrows), scale bar = 50 um, (A’’) stage II follicle, scale bar = 25 um. (B) stage V follicle, scale bar = 200um. (B’) higher magnification illustrates basement membrane surrounding the follicle (arrows) and basement membrane within the theca cell layer (th) (asterisks), scale bar = 25 um. (C) stage VI follicle, scale bar = 200um. (C’) double basement membrane around the granulosa cell layer (g) remains, scale bar = 50 um. (D) stage VIII follicle, scale bar = 200um. (D’) double basement membrane around the granulosa cell layer remains, scale bar = 50 um. (E) stage IX follicle, scale bar = 200um. (E’/E’’) higher maginification of theca cell layer, only one thin basement membrane detected, scale bar = 25 um. **F-H.** Immunofluorescence staining of follicle development. DAPI (cyan) and WGA (magenta) staining. Left maximum intensity projection, scale bar = 200 um. middle 3 columns individual z-sections, scale bar = 100 um. Boxes indicate higher magnification in the right column, scale bar = 50 um. (F) germinal bed, z85 asterisks highlight germ cells, pf = primordial follicles. z100/z110 dashed lines indicate the border of the germinal bed (gb) to the left, maturing follicles (mf) to the right. z110 higher magnification asterisks indicate stage II. (G) stage II IV follicles. higher magnification asterisks highlight double basement membrane. (H) stage III-V follicles. Arrows indicate outer basement membrane deposited pericellularly. Right: higher magnification of perforations in inner basement membrane.

Once the granulosa and theca layers are fully established at stage V, two layers of basement membrane can be detected (Figure 9B), with one thin layer directly overlaying the maturing follicle (asterisks), while the other basement membrane is located at the granulosa-theca interface and is much thicker (arrows) (Figure 5B’). This double membrane on either side of the granulosa cell layer is retained throughout follicle maturation (Figure 9C/C’/D/D’). At stage IX, when the granulosa cell layer is lost (Figure 9E), only the thin basement membrane layer directly overlying the oocyte remains (Figure 9E’/E’’).

To investigate basement membrane localization in 3D, we performed whole-mount confocal imaging of ovaries containing stage I-VIII follicles, stained with Wheat Germ Agglutinin (WGA, a lectin that binds proteins abundant in basement membranes). The germinal bed was localized based on its position and morphology (Figure 9F z85/z100), and the primordial follicles were identified as small pockets in tightly packed nuclei containing one slightly enlarged nucleus (Figure 9F z85 DAPI). Interestingly, these individual primordial follicles could also be distinguished with WGA staining, which formed distinct pockets around each oogonial cell (Figure 9F z85 WGA). The germinal bed appears separate from the follicles undergoing maturation (Figure 9F z100/z110, dashed line). At stage II (Figure 9F z110, higher magnification, asterisks), two basement membranes can be observed on either side of the granulosa cell layer. This double layer then becomes more pronounced during subsequent stages (Figure 9G). In older follicles (stages III-VII, Figure 9H), the inner basement membrane exhibits perforations (Figure 9H z140 higher magnification, right column). In contrast, the outer basement membrane instead appears to be deposited not only on the basal side of the cells, but also in-between the cells, enabling the definition of cell shapes (Figure 9H, WGA arrows). Thus, distinct basement membrane distribution patterns accompany follicle maturation, and include a characteristic pattern in the geminal bed and the acquisition of a double membrane around the granulosa cells as evidenced via both histology staining and WGA confocal imaging.

## Discussion

In this study, we provide a comprehensive description of ovarian oocyte and follicle development in the brown anole. Using brightfield and CT imaging as well as SEM, histological staining, and confocal microscopy, we define 10 consecutive stages of oogenesis and folliculogenesis. Our first 6 stages are combined into stage 1 of the staging system described for *Anolis pulchellus* (Ortiz & Morales, 1974) and we defined one novel stage that precedes the 9 stages defined for *Tropidurus* (da Silva et al., 2018). To enable cross-species comparison, we designated these stages Ia and Ib instead of creating a novel competing staging system. Although ovarian follicles and follicle development have been described in the genus *Anolis* before (Laughran et al., 1981; Ortiz & Morales, 1974), our study contributes substantial new detail by providing a more detailed description dividing oogenesis not into four, but into 10 stages based on granulosa cell morphology, yolk development, and follicle size changes in the brown anole ovary.

We found that in the brown anole ovary, the follicular epithelium undergoes stereotyped changes in cell type composition, and we hypothesize that each cell type has a different functional role in support of follicular development. We observed changes in the ratio of granulosa cell types as follicles mature, consistent with patterns observed in other lizards, and we identified the three known granulosa cell types: small cells, intermediate cells, and pyriform granulosa cells. While granulosa cells are known to play a critical role in the development of the mouse ovary (Nilsson & Skinner, 2001), we currently only have a limited understanding of the role of the different granulosa cell types in the brown anole ovary during oocyte development. Previous analyses of reptile ovaries posited different hypotheses regarding the function and origin of the different granulosa cell types (Filosa et al., 1979; Klosterman, 1983; Neaves, 1971). However, no functional studies have tested these hypotheses to better understand the mechanism governing the consecutive reorganization of this highly dynamic structure. With our work, we lay the foundation for such stage-specific future functional analyses.

The exponential increase in follicle volume observed in our dataset is likely driven by the accumulation of yolk in the oocyte. We observe discernable stages of yolk deposition within the follicles but could not link the patterns to specific features within the granulosa cell layers. We observed that oogonia in the germinal bed do not stain positive for eosin, indicating a lack of cytoplasmic protein (Nuovo, 2020). Later stages of follicles had eosin-positive cytoplasmic material, which is indicative of protein deposition. The protein required for yolk development, vitellogenin, has been reported in turtles to be transported through blood vessels and is deposited into the oocyte (Gapp et al., 1979; Ho et al., 1982). Interestingly, we observed perforations of the basement membrane in vitellogenic follicles that could enable growth of the follicle but may also serve as sites of yolk transfer through the basement membrane. Further studies are required to understand the precise mechanism of yolk deposition in the brown anole oocyte.

We observe that a basement membrane is established in young follicles and that at older stages of follicular development, a double basement membrane is present. Basement membranes are assembled from laminin secreted by epithelial cells and are required to establish appropriate tissue architecture and structure (Ecay & Valentich, 1992). In our data, we show that gaps develop in the basement membrane of mature, pre-ovulatory follicles. In mice, the embryonic basement membrane has been shown to be actively remodeled during implantation and gastrulation so that perforations appear and are distributed asymmetrically, contributing to anterior-posterior axis formation (Chen et al., 2025; Kyprianou et al., 2020). The perforations we observe in the brown anole oocyte basement membrane surrounding the oocyte could be part of an active remodeling process during oocyte maturation. These perforations could also serve to facilitate vitellogenin acquisition from the bloodstream, as discussed above. The double basement membrane is also remarkable. A dual membrane is characteristic of other organs, for instance, the glomerular basement membrane in the developing kidney. In this case, the dual membrane results from fusion of membranes secreted by epithelial and endothelial cells (Abrahamson, 1987). In our case, it is unclear exactly what the origin of the dual membrane is. Additional work on the development and function of the oocyte basement membrane would therefore be beneficial.

We identified a single germinal bed per ovary which is separated from the maturing follicles of the ovary. Mechanisms leading to oogenesis initiation, including how maturing follicles leave the germinal bed to begin maturation are poorly understood. This movement and rearrangement of the basement membrane may be orchestrated by granulosa cells. However, we still need to gain a deeper understanding of the gene expression patterns of the cells that constitute the germinal bed as well as the early follicles to answer this question. Similarly, because other non-avian reptile species, such as lizards in the genera *Sceloporus* and *Basiliscus* (Jones et al., 1982), can have multiple germinal beds per ovary and because clutch size is highly variable across reptiles, examining how the number of oogonial divisions per reproductive cycle correlates with clutch sizes would be interesting.

Our study establishes that brown anole oogenesis occurs in a pattern conserved with other lizards. We present a staging table of follicle development that will lay the basis for functional and cross-clade comparative studies of oogenesis, further cementing the brown anole as a representative species of the genus *Anolis*. Through this precise staging system, the timing of genome editing may be more precisely defined enabling reproducibility and identification of the optimal timepoint for genome modification. Understanding oogenesis in reptiles also has a broader potential to influence the field of Assisted Reproductive Technology. Identification of the factors that facilitate continuous germ cell production within the germinal bed across the adult life of a non-avian reptile may open novel avenues to counter infertility in mammals that have a finite number of oocytes (Hutchinson et al., 2025).

## Contributions

BKK & AW conceived of the project, carried out, and analyzed the experiments. EJV carried out digital segmentation and volumetric analysis of CT datasets. NAS helped with experiment planning. KS, TJS, BKK and AW collected *A. sagrei* females. ZBG carried out histological analysis. HW carried out histology. MMC carried out STEM. FH, SAW, TJS, RRB and PAT supervised the project and provided critical feedback. BKK & AW wrote the manuscript with the help of SAW, RRB, TJS, NAS, FH, and PAT.

## Acknowledgements

The authors thank all members of the Trainor, Behringer & Williams labs for critical feedback. We thank Edward Stanley and Martin Cohn at the University of Florida for support generating the CT data. Research in the Trainor lab is funded by the Stowers Institute for Medical Research (1008). The Behringer lab is supported by NIH grants 5R01HD113569 and HD30284 you and the Ben F. Love Endowment to Richard R. Behringer. Bonnie K. Kircher was supported by National Science Foundation (NSF) Postdoctoral Research Fellowship in Biology (PRFB), Division of Biological Infrastructure (DBI) 2209150 and NIH T32HD098068. MicroCT scanning was supported by a NSF Graduate Research Fellowship awarded to Bonnie K. Kircher. Veterinary services were supported by NIH grant CA16672. The Sanger lab is funded through the USA National Science Foundation (#1942250). A.W. is the recipient of an independent postdoctoral research fellowship of All Souls College, University of Oxford, and received a travel grant from the Cambridge Philosophical Society. A.W. and F.H. are supported by the Newton Trust, the European Research Council (695669) and the Engineering and Physical Sciences Research Council (EP/Y032756/1). NAS is supported by a K99/R00 Pathway to Independence award from the National Institute of Child Health and Human Development (HD114881).

## Methods

### Data Availability

All original data underlying this manuscript can be accessed upon publication at the Stowers Original Data Repository at https://www.stowers.org/research/publications/libpb-XXXX CT data for anoles 1 and 2 reanalyzed from Kircher and McCowen et al. (Kircher, McCown et al. 2024). All scans have been uploaded to Morphosource (Anole 1 Morphosource ID: 000590186, Anole 2 Morphosource ID: 000608703, Anole 3 Morphosource ID: 000702213).

### Collection of females

All animal collection and procedures used in this study were approved by MD Anderson Cancer Center IACUC (#00002169), the University of Florida IACUC (# 201408266) or the Loyola University Chicago IACUC (#3662). *A. sagrei* females were collected at the Houston Arboretum in summer 2021, Gainesville, FL, summer 2016-17, and in Miami, FL, May 2023 according to state regulations. Females were euthanized according to one of the following protocols. In Houston, females were anesthetized with 1% Tricaine followed by 50% tricaine injection. In Gainesville, anoles were euthanized using Euthasol, fixed using 10% formalin. In Miami, females were placed in a cooling box with an ice pack to slow down metabolism for anesthesia and then euthanized through decapitation. In Houston, anoles were anesthetized by 1% Tricaine does by weight and then euthanized by a 50% tricaine injection dosed by weight (Conroy 2009).

### MicroCT

*Anolis sagrei* were fixed using 10% formalin and stored in 70% ethanol. Then, specimens were stained with 1.75% aqueous Lugol’s iodine (Gignac, Kley et al. 2016) and their whole body was visualized with X-ray computed tomography. All samples were scanned on GE V|Tome|X M 240 system at Nanoscale Research Facility at the University of Florida. Source and detector settings are listed in Table 1. Tomograms were generated using Reconstructor software v.16.0.11592 (Carl Zeiss Microscopy, GmbH) and segmentation was done using 3D-slicer (Fedorov, Beichel et al. 2012). All animal collection and procedures for these methods were approved by the University of Florida IACUC.

### Histology

Samples were previously fixed and stored in 70% ethanol as described above. Samples were paraffin processed (Milestone, Pathos Delta Microwave Tissue Processor) and subsequently embedded in paraffin wax (Cancer Diagnostics, PureAffin® R56). Samples were then sectioned at a thickness of 5 µm on a microtome (Leica RM2255). H&E staining was performed using an automatic stainer (DP360, Dakewe (Shenzen) Medical Equipment Co.) with ST Infinity H&E Reagents (Leica Biosystems Cat. 3801698). Slides were mounted with Cytoseal 60 mounting media (VWR, 48212-187).

### Electron microscopy

Samples were prepared for electron microscopy as previously described (Weberling et al., 2025). Sections were cut at 80nm with a Diatome diamond knife and stained with 6 minutes each of 4% uranyl acetate in 70% methanol and Sato’s triple lead stain before imaging in a Zeiss Merlin SEM with aSTEM detector at 26kV and 700pA.

### Confocal imaging

Ovaries were incubated in DAPI and Wheat Germ Agglutinin (AF-594) and imaged on a Zeiss LSM800 or Zeiss LSM900. Tile scans were collected with 15% overlap.

### Computational Analysis

Brightfield images were rendered using the Helicon Software.

All quantitative analyses were carried out using FIJI (Schindelin et al., 2012).

### Quantitative analysis

#### Volumetric analysis of CT data

We reanalyzed the datasets published in Kircher and McCowen et al. (Kircher, McCown et al. 2024) to represent anoles 1 and 2 Morphosource ID: 000590186, 000608703). Anole 3 has been uploaded to Morphosource (ID: 000702213). To compare the volume of each follicle on the ovary, we digitally segmented in 3D slicer (Fedorov et al., 2012) each follicle and determined the volume from the digital 3D dataset. We measured the snout-vent-length (SVL) of the lizard and divided the ovarian follicle volume by the SVL of the lizard to correct for body size. We log-transformed the body-size-corrected follicle volume and multiplied that volume by -1. We also ranked each follicle by the volume of the follicle compared to the others on the ovary. The smallest follicle is ranked number 1 and each follicle with increasing volume has a progressively higher rank. We graphed the body-size-corrected-volume against follicle rank and added error bars representing standard deviation from the mean value of each follicle rank using R (version 4.5.0) in R Studio (2025.05.0+496). We also analyzed the change in follicular volume across each individual ovary sampled in our dataset using a linear regression model in R (version 4.5.0) in R Studio (2025.05.0+496).

#### Percentage of large cells

For each follicle the cross section of the largest diameter was chosen. All cells within the granulosa cell layer were counted. The number of large cells (including intermediate and pyriform cells) was divided by the total number of cells. The percentage was plotted using R (version 4.5.0) in R Studio (2025.05.0+496). We fit a linear regression model using the geom_smooth function in ggplot2.

#### Follicular epithelium width

Linear measurements of the follicular epithelium were taken for each follicle again at the cross section that showed the largest diameter for the respective follicle. Width measurements were collected in three places around the circumference of the follicle. The average of these three measurements was plotted against the follicle stage using R (version 4.5.0) in R Studio (2025.05.0+496). A generalized additive model was fitted using the geom_smooth function in ggplot2.

#### Plot profiles

Plot profiles were drawn across the width of the oocyte, excluding the follicular epithelium. Plot profiles were drawn to avoid the nucleus and any sectioning artifacts (tears, wrinkles, etc.). Line width of the plot profile was 10 uM for pre-vitellogenic follicles and 100 uM for early vitellogenic, vitellogenic, and mature vitellogenic follicles. Higher mean grey value indicates higher density while lower mean grey value indicates lower density.

