## Supplementary material for "Oogenesis and germinal bed morphology of the brown anole (*A. sagrei*)": tables

Table 1. Regression statistics for follicular volume across six individual ovaries analyzed.

| Anole | Ovary | p-value | F(1,3) | R-squared |
|---|---|---|---|---|
| 1 | L | 0.002884 | 81.22 | 0.9525 |
| 1 | R | 0.0001403 | 625.2 | 0.9936 |
| 2 | L | 0.003535 | 70.62 | 0.9457 |
| 2 | R | 0.01361 | 27.34 | 0.8682 |
| 3 | L | 0.004972 | 55.77 | 0.9319 |
| 3 | R | 0.003753 | 67.76 | 0.9435 |

Table 2.

| ID | Color | Yolk Stage | Follicle Number | Follicle Stage | Small Cell Count | Large Cell Count | Total | Small% | Large% | Follicle Area um | Follicle Area um | Follicle Width 1 | Follicle Width 2 | Follicle Width 3 | Average Follicle Width |
|---|---|---|---|---|---|---|---|---|---|---|---|---|---|---|---|
| M122C | Blue | pv | 1 | 2 | 22 | 0 | 22 | 100 | 0 | 1002.667 | 1.002667 | 2.828 | 1.795 | 3.145 | 2.47 |
| M127C | Red | pv | 7 | 2 | 30 | 5 | 35 | 85.7142857 | 14.2857143 | 1492 | 1.492 | 5.27 | 4.269 | 4.955 | 4.83133333 |
| M122C | Blue | pv | 2 | 3 | 32 | 5 | 37 | 86.4864865 | 13.5135135 | 1730.444 | 1.730444 | 5.088 | 4.346 | 7.032 | 5.48866667 |
| M127C | Red | pv | 6 | 3 | 55 | 27 | 82 | 67.0731707 | 32.9268293 | 6282.667 | 6.282667 | 13 | 12.737 | 11.101 | 12.2793333 |
| M121D | Black | pv | 2 | 4 | 58 | 64 | 122 | 47.5409836 | 52.4590164 | 15193.22 | 15.19322 | 12.089 | 7.769 | 9.22 | 9.69266667 |
| M121D | Black | pv | 4 | 4 | 88 | 75 | 163 | 53.9877301 | 46.0122699 | 11097.889 | 11.097889 | 8.969 | 13.876 | 9.597 | 10.814 |
| M121D | Black | pv | 3 | 4 | 60 | 158 | 218 | 27.5229358 | 72.4770642 | 33913.778 | 33.913778 | 12.512 | 13.271 | 12.944 | 12.909 |
| M122C | Blue | pv | 3 | 4 | 73 | 40 | 113 | 64.6017699 | 35.3982301 | 16314.778 | 16.314778 | 14.461 | 11.324 | 10.05 | 11.945 |
| M122C | Blue | pv | 4 | 4 | 79 | 62 | 141 | 56.0283688 | 43.9716312 | 27410.111 | 27.410111 | 15.202 | 12.293 | 13.671 | 13.722 |
| M127C | Red | pv | 5 | 4 | 85 | 85 | 170 | 50 | 50 | 12911.778 | 12.911778 | 13.864 | 16.035 | 16.974 | 15.6243333 |
| M127C | Red | pv | 8 | 4 | 66 | 124 | 190 | 34.7368421 | 65.2631579 | 24580.222 | 24.580222 | 17.714 | 16.364 | 18.342 | 17.4733333 |
| M121D | Black | ev | 6 | 5 | 171 | 306 | 477 | 35.8490566 | 64.1509434 | 63410.899 | 63.410899 | 17.537 | 14.941 | 18.028 | 16.8353333 |
| M122C | Blue | pv | 5 | 5 | 177 | 182 | 359 | 49.3036212 | 50.6963788 | 162134 | 162.134 | 18.487 | 24.413 | 28.674 | 23.858 |
| M122C | Blue | pv | 7 | 5 | 139 | 375 | 514 | 27.0428016 | 72.9571984 | 266142.444 | 266.142444 | 32.77 | 42.165 | 48.671 | 41.202 |
| M127C | Red | pv | 9 | 5 | 35 | 252 | 287 | 12.195122 | 87.8048781 | 43227 | 43.227 | 22.01 | 18.273 | 18.812 | 19.6983333 |
| M127C | Red | ev | 4 | 5 | 114 | 200 | 314 | 36.3057325 | 63.6942675 | 58554 | 58.554 | 15.563 | 21.715 | 19.128 | 18.802 |
| M121D | Black | ev | 5 | 6 | 80 | 580 | 660 | 12.1212121 | 87.8787879 | 200832 | 200.832 | 17.92 | 14.981 | 18.523 | 17.1413333 |
| M121D | Black | ev | 7 | 6 | 71 | 693 | 764 | 9.29319372 | 90.7068063 | 129527.667 | 129.527667 | 19.047 | 19.531 | 29.494 | 22.6906667 |
| M122C | Blue | ev | 6 | 6 | 109 | 176 | 285 | 38.245614 | 61.754386 | 121226.444 | 121.226444 | 21.797 | 19.821 | 26.874 | 22.8306667 |

| | | | | | | | | | | | | | | | |
|---|---|---|---|---|---|---|---|---|---|---|---|---|---|---|---|
| M122C | Blue | v | 8 | 6 | 30 | 548 | 578 | 5.19031142 | 94.8096886 | 644370.889 | 644.370889 | 30.806 | 27.014 | 24.775 | 27.5316667 |
| M127C | Red | v | 3 | 6 | 148 | 484 | 632 | 23.4177215 | 76.5822785 | 188771 | 188.771 | 22.103 | 20.276 | 21.095 | 21.158 |
| M127C | Red | ev | 2 | 6 | 171 | 373 | 544 | 31.4338235 | 68.5661765 | 99194 | 99.194 | 23.286 | 18.236 | 19.235 | 20.2523333 |
| M127C | Red | v | 1 | 7 | 54 | 617 | 671 | 8.04769002 | 91.95231 | 289749.333 | 289.749333 | 11.353 | 16.09 | 13.017 | 13.4866667 |
| M121D | Black | V | 1 | 8 | 103 | 529 | 632 | 16.2974684 | 83.7025317 | 318553.222 | 318.553222 | 10.833 | 11.319 | 11.108 | 11.0866667 |
| M121D | Black | mv | 8 | 9 | N/A | N/A | N/A | N/A | N/A | N/A | N/A | 4.069 | 3.606 | 4.738 | 4.13766667 |
| M122C | Blue | mv | 9 | 9 | N/A | N/A | N/A | N/A | N/A | N/A | N/A | 8.028 | 4.384 | 4.679 | 5.697 |
| M127C | Red | mv | 10 | 9 | N/A | N/A | N/A | N/A | N/A | N/A | N/A | 5.09 | 6.037 | 6.083 | 5.73666667 |
